# Pairwise Causal Discovery in Biochemical Network: A Survey on Directionality Inference within Complex Networks from Stationary Observations

**DOI:** 10.1101/2025.07.22.666141

**Authors:** Nava Leibovich, Miroslava Cuperlovic-Culf

**Affiliations:** National Research Council of Canada, NRC-Fields Mathematical Sciences Collaboration Centre, 222 College street, Toronto, ON, M5T 3J1, Canada; Digital Technologies Research Centre, Bldg M50, National Research Council Canada, 1200 Montreal Road, Ottawa, ON, K1A 0R6, Canada; Department of Biochemistry, Microbiology and Immunology, Faculty of Medicine University of Ottawa, 451 Smyth Road, Ottawa, ON, K1H 8M5, Canada; Ottawa Institute of Systems Biology, University of Ottawa, 451 Smyth Road, Ottawa, ON, K1H 8M5, Canada

## Abstract

Metabolic networks map complex biochemical reactions within organisms, which is crucial for understanding cellular processes and metabolite flow. This study focuses on inferring the directionality of interactions in metabolomics networks. Given the challenge of using steady-state data, we benchmark various methods, including statistical scores and neural network approaches, on synthetic yet realistic biological models. Our findings highlight the relative success of a few methods in some cases where the interaction mechanism is known, whereas other methods show limited effectiveness.

## 1. Introduction

Metabolic networks function as intricate maps of biochemical reactions within an organism, illustrating the complex web of interactions that govern cellular processes. Understanding the directionality of these reactions is crucial for tracking the flow of metabolites through pathways and for illuminating the functional dynamics of cellular metabolism. This information is relevant across various fields, including systems biology, biotechnology, and drug discovery [1–5]. In this context, we concentrate on the interpretation of the metabolomics network.

The metabolome consists of hundreds of correlated compounds, offering substantial information within the metabolomics network that extends beyond the mere levels of individual metabolites [6–8]. Focusing solely on individual metabolites can obscure potential intervention targets due to issues with collinearity and confounding factors; however, network models can effectively incorporate these interrelationships within the metabolome [6–10]. The significance of biological networks in human diseases has been widely acknowledged [11, 12], particularly in their role in identifying causal associations [13].

Interaction networks are typically inferred using time series measurements or pseudo-time trajectories, employing statistical tools such as Bayesian inference and maximum likelihood alongside machine learning algorithms [14–24]. Notably, a wide range of recent studies have explored methods for inferring networks from temporal data, as highlighted in the partial list as follows [25–35]. However, it is important to note that these methods require recorded synchronized ordered data, which is often unmeasurable in various observational contexts.

Here we aim to infer the direction of interactions from stationary data associated with the analyzed dataset, see Fig. 1. In studies of biochemical reaction networks, one can derive the *functional* connectivity, which provides valuable insights into the statistical dependencies arising from the collective dynamics of interactions between pairs of units [36–42]. These statistical dependencies among components can be characterized by well-known measures such as correlation, mutual information, or their analyses using silencing methods [40–44]. However, these measures are symmetric and consequently do not capture the direction of interactions, resulting in the inference of non-directed networks. In contrast, non-symmetric metrics, such as partial correlations and dependency analyses, necessitate recorded data from all other interacting variables within the network [45–49]. Such comprehensive data is often unavailable in many systems.

**Figure 1:**
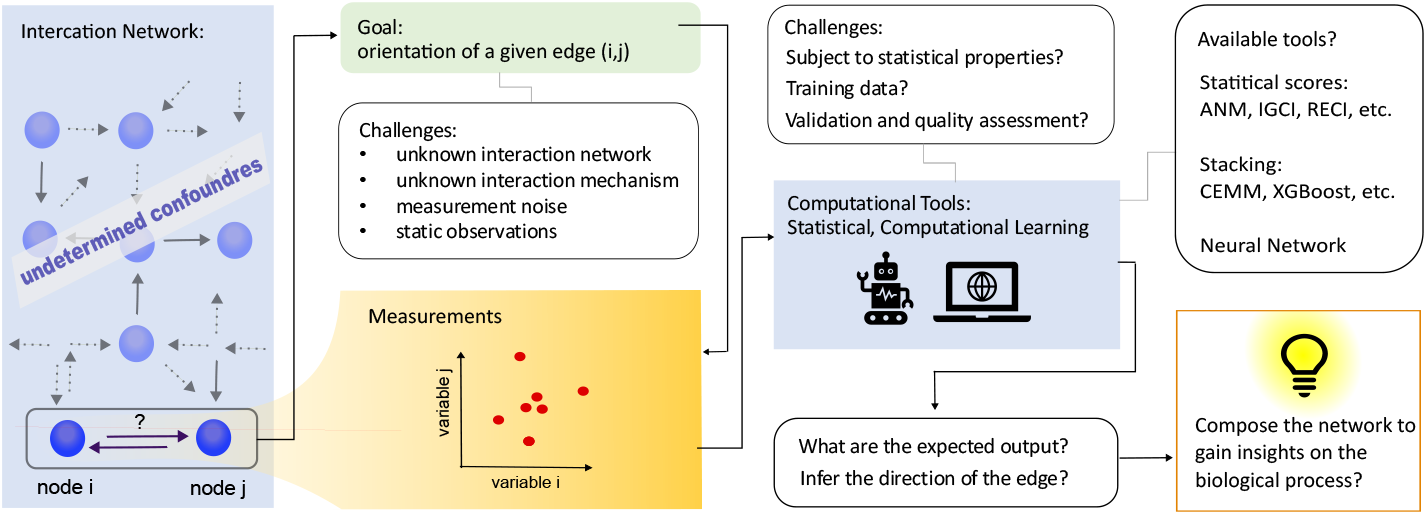
The main goal is to infer the direction of interaction between two variables within a metabolic network. Our focus is on methods designed to uncover the directional influence between these two variables. However, determining the orientation of the network from static data poses a great challenge, necessitating keen analytical and computational techniques.

In particular, we recognize that the complete metabolic network under consideration has not been fully characterized. As a result, neither the overall network topology nor other confounding variables have been adequately described or measured. Consequently, we focus on methods designed to infer the pairwise causal direction between two correlated variables. Furthermore, we emphasize that earlier benchmark analyzes of pairwise inference methods [50–52] evaluate performance with respect to various causal mechanisms between pairs of variables, without taking into account the context of interaction networks in biological processes. Such networks can introduce intricate influences due to signal propagation, even when the two variables do not directly interact with one another.

Determining the network orientation from steady-state data is therefore challenging. First, we examine various methods for deciphering interaction directions using steady-state data and highlight their domains of failure and success. We survey both statistical scores which allow inference from small data sets, and methods based on machine learning and neural network approaches which require large data sets for training the models. We tested these methods on multiple interaction mechanisms, with and without additional noise.

## 2. Main Results

As mentioned, we aim to infer the direction of interaction, within a partially observed interaction network between biochemical components, to gain insights into some biological processes. To do so, we benchmark existing methods to infer directionality from pairwise data. Our contribution lies in the following. Previous comparisons between methods collate various interaction mechanisms, combining linear and non-linear effects, and additive and multiplicative noise. Nevertheless, these previously examined mechanisms do not necessarily capture the complexity of biochemical reaction networks; which possess multiple confounders that may remain unknown, including biochemical cycles and interaction loops, complex signal propagation throughout the network, and which the pairwise interactions mechanism is unknown.

### 2.1. Benchmarking Methods for Analyzing Synthetic Data

We use several interaction mechanisms within complex networks. Specifically, we use the Michaelis-Menten model, which describes gene interaction networks [53], as well as networks of coupled Rössler oscillators, and coupled Goodwin oscillators which represent a prototypical biological oscillator that characterizes various biological processes such as circadian clocks [54, 55]. Further details on these models can be found in Appendix A. The interaction networks are random graphs following the Erdös–Rényi model. In Fig. 2 we show the observations of pairs for the interaction mechanisms considered. Given that both variables are affected by additional influences from other network variables, along with inherent noise for each variable which also propagates throughout the network, determining the direction of the interaction is highly challenging.

**Figure 2:**
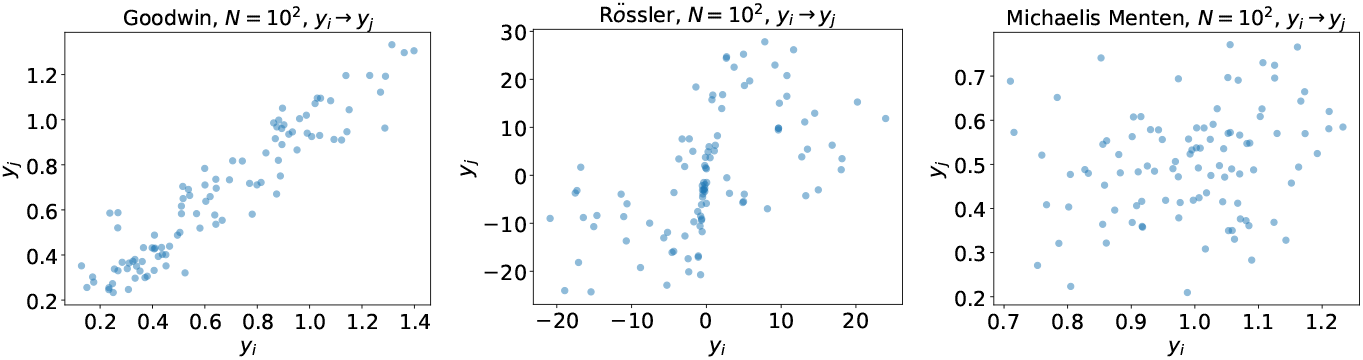
Realizations of interacting variables’ pairs (*y*_*i*_, *y*_*j*_) for different interaction mechanisms; Good-win, Rössler, and Michaelis-Menten models (left to right correspondingly). For consistent visualization, we present the direction of interaction *y*_*i*_ *→ y*_*j*_ for all panels, where for each we sample *N* = 10^2^ data points.

We investigated a range of methods, including those grounded in statistical scores, stacking techniques, and artificial neural networks (NNs). The statistical scores we analyzed include the Additive Noise Model (ANM), Conditional Distribution Similarity (CDS), Information Geometric Causal Inference (IGCI), and Regression-Error-based Causal Inference (RECI). Additionally, we examined various stacking methods, such as voting, eXtreme Gradient Boosting (XGBoost), and a recently introduced Causal Ensemble Learning method based on Support Measure Machines (CEMM). Furthermore, we explored several proposed deep neural network architectures. A comprehensive overview and description of these inference methods can be found in Table 1 and in Appendix B. The implementations were based on [56] wherever applicable.

**Table 1:**
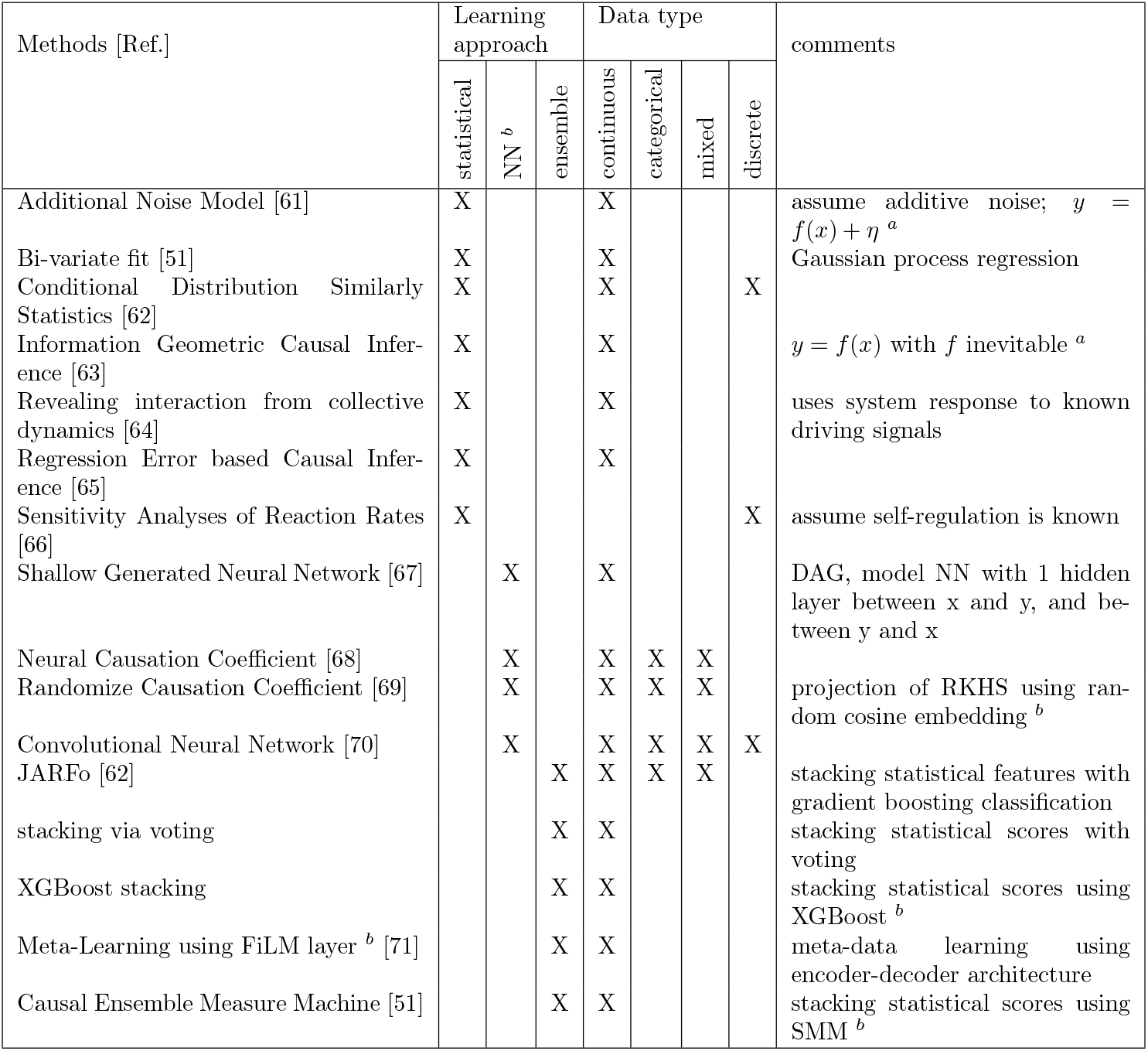
Pairwise interaction orientation methods. ^*a*^ For uniform notation we assume that the variable *x* affects the variable *y*, i.e. *x → y*. The noise term is marked with *η*. ^*b*^ Abbreviations: NN = neural networks, RHHS = reproducing kernel Hilbert space, DAG = directed acyclic graph, FiLM = Feature-wise Linear Modulation, XGBoost = eXtreme Gradient Boosting, SMM = Support Measure Machine

The techniques that have been tested provide a rich composition of methods, each incorporating different approaches to inference. This variety offers a solid foundation for benchmarking assessment. However, we have not examined all possible methods that have been proposed in the literature. Certain methods were excluded for several reasons, including limitations in computational capacity, the unavailability of published code from the relevant articles, and a lack of applicability to our data. Specifically, some methods require information that is inaccessible or involve data types that do not align with the data of our interest. As mentioned, the methods are reviewed in Table 1 and Appendix B.

#### 2.1.1. Performance on Random Synthetic Networks

We show in Fig. 3 that for the biochemical reaction networks we examined, most methods we assessed struggle with direction determination, as no method presents excellent accuracy, say beyond 0.9. This somewhat inferior accuracy is expected, especially for the statistical scores, since they rely on strong assumptions about the statistical characteristics of the data, which are not necessarily complied within our systems. In general, accuracy around 0.5 indicates a random choice between two equally probable cases, while accuracy lower than that implies systematic errors yielded in the statistical scores from their unfulfilled assumptions.

**Figure 3:**
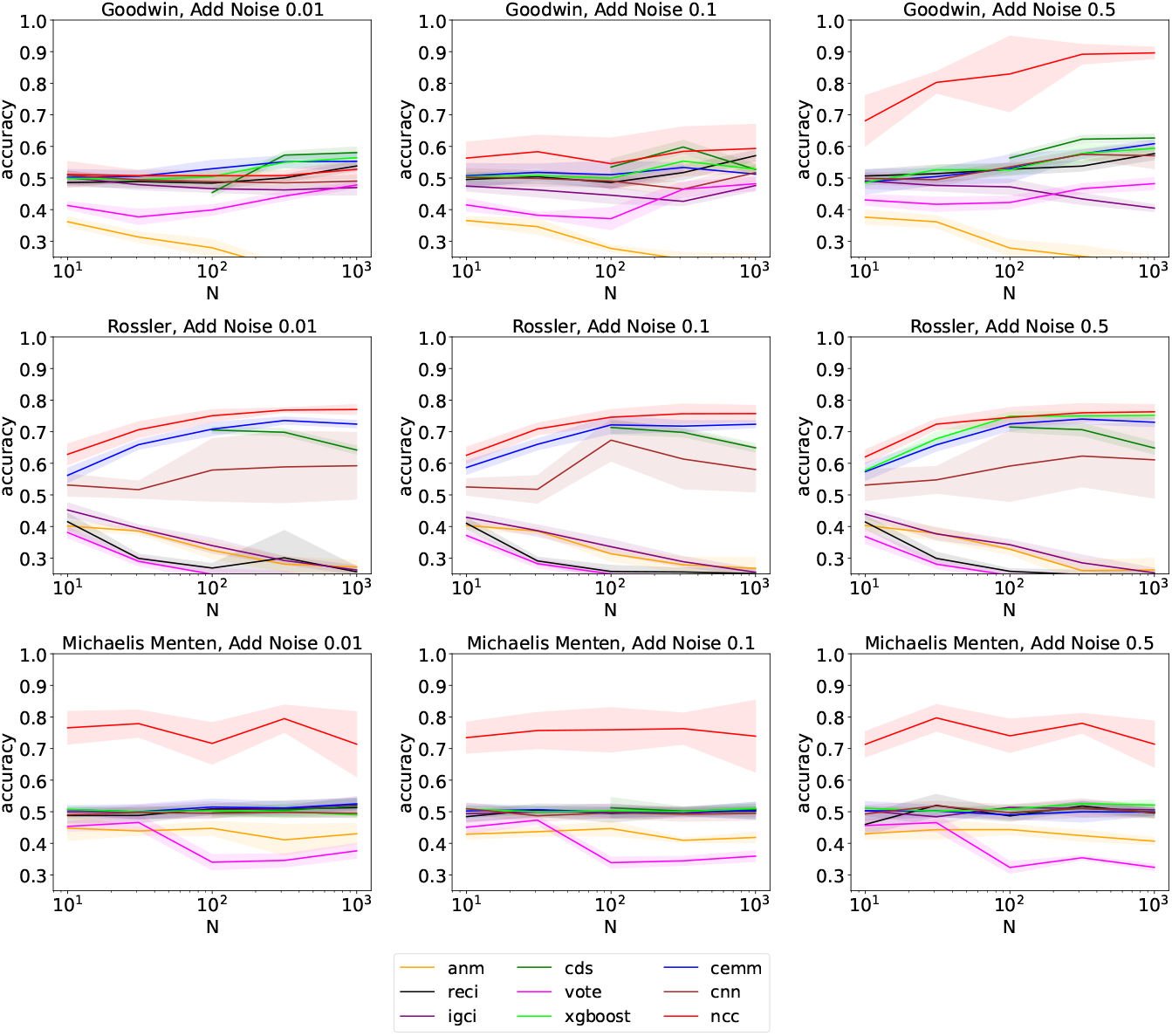
Results for the tested directionality inference methods for various interaction mechanisms.

However, we found that some methods perform better than others. In particular, the NCC method yields results that surpass the baseline of random choice located at 0.5, with accuracies ranging between 0.7 to 0.9. We find that the NCC results exceed other methods for various interaction mechanisms and noise scales that were considered. We further investigate insights from these results in the next subsections.

Interestingly, for the Rössler coupled oscillators, we found that the CEMM, XG-Boost, and CDS methods produced relatively successful results, albeit falling short when compared to the NCC method (see Fig. 3). Among the stacking methods, XGBoost and CEMM demonstrated accuracy that places them secondary to NCC, achieving an approximate accuracy of 0.7. Additionally, for the Rössler coupled oscillators, the statistical score from CDS exhibits similar accuracy for larger values of *N*. In contrast, for the other interaction mechanisms we examined, the CDS, CEMM, and XGBoost stacking methods, along with the other inference approaches tested—excluding NCC—failed to accurately infer the direction of the interaction.

Moreover, our findings indicate some dependence on the noise level. We tested the performance for three noise levels. For data synthesized with the Goodwin mechanism, a higher noise level seems preferable, and success was found for the NCC method. Other dynamic mechanisms show comparable results for all noise levels tested. Additionally, the finite number of sampled data points, *N*, induced an additional measurement noise from the finite sampling itself. For small *N*, most methods show accuracy close to 0.5, which signifies the chance accuracy of a dummy classifier for the two possible directions. As *N* increases, some methods improve their accuracy, while for others the accuracy stays at the random chance or even decreases below it, due to the systematic errors that emerged from the statistical assumption. Given that observations are typically in the size of 10^2^ − 10^3^ samples, we do not consider a larger set of data points.

#### 2.1.2. Further inspection of the NCC model

To expand the analyses even further, we have tested the NCC method as a candidate for a directionality inference method. We first examine the performance of the model trained on a given dataset drawn from one mechanism, to infer interaction direction from another mechanism, see Fig. 4. As expected, each NCC model performs better on the data mechanism it was trained on. Obviously, for a trained model to be effective, the training data must be similar to the data aimed to infer. Nevertheless, in the biological processes under question, one does not necessarily have such information.

**Figure 4:**
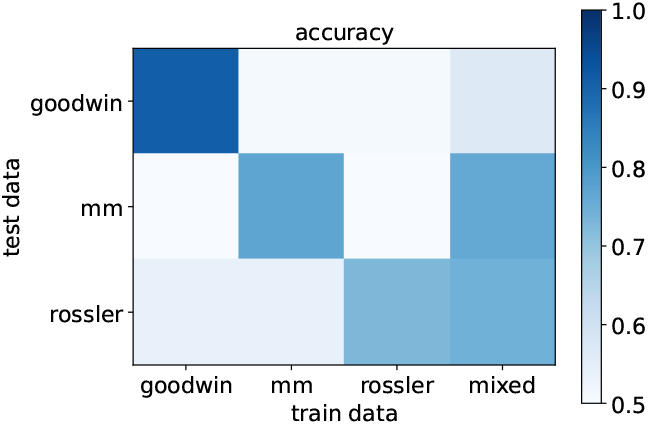
Testing data vs trained NCC models. As expected, using training and testing datasets from different mechanisms is ineffective.

We also trained an additional model using simulated data generated with randomly selected interaction mechanisms, referred to as ‘mixed’. Unlike before, we did not specify particular mechanisms. This new model yielded slightly improved results, exceeding the random accuracy for two different mechanisms we examined.

The analysis also considers the correlation coefficients of Kendall’s *τ* and Spearman’s *ρ*, along with the mutual information between the two variables. These measures evaluate the observed dependence between the two variables in question. Higher values suggest that the dynamics of one variable depend on the other. We found that a strong dependence between the variables results in a higher probability of successfully determining the interaction direction, see Fig. 5. In our synthetic data, all examined edges represent defined interactions between the nodes, where one variable categorically influences the other. However, weak interactions may be empirically detected due to sampling errors, the propagation of noise from other parts of the network, and the influence of various confounding factors on the dynamics of both variables.

**Figure 5:**
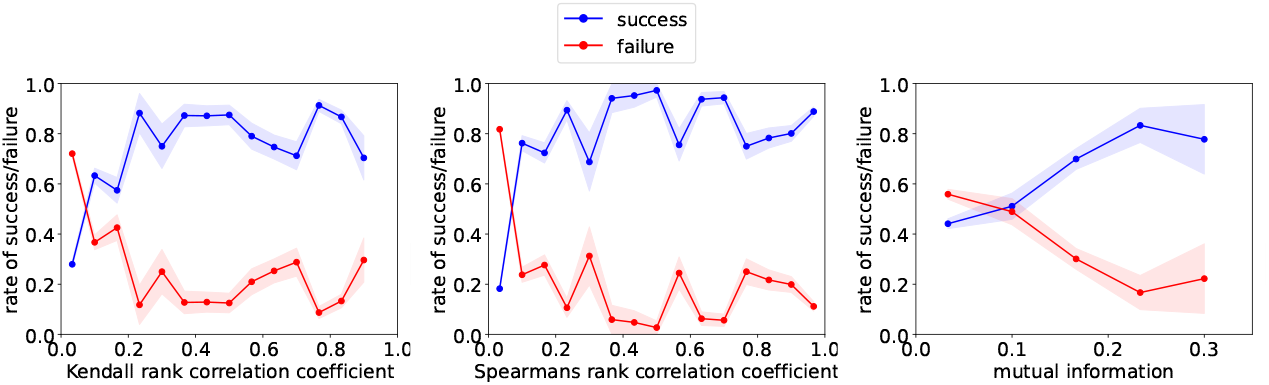
Dependence quantities influence the probability for meaningful training of the NCC model and thus to a successful inference. Results for the Kendall *τ* coefficient (left), Spearman *ρ* coefficient (middle), and mutual information (right) show that stronger dependence between the two variables corresponds to increased success rates in determining the direction of interaction between them.

#### 2.1.3. Model Performance for Realistic Biological Systems

We test the applicability of the above models for inference of the interaction direction within realistic biological processes. To do so, we simulate these biological processes and test the performance of the inferred interaction direction. Specifically, we first use a previously proposed biologically realistic system - a model that is based on the *E. coli* gene regulatory network (GNR) provided in the DREAM challenge [57–59]. In addition, we simulated the Bile Acid Synthesis Pathway (BASP) model specified in [60] for describing part of the cholesterol metabolism. Finally, we simulate sphyngolipid metabolism using computationally determined enzyme kinetic parameters and a detailed metabolic network based on KEGG: map00600. The details of all models are provided in Appendix A.

Testing the models on synthetic yet realistic biological process data enables us to evaluate the anticipated performance of the inference method in a controlled environment before applying it to actual data of interest. We utilize our pre-trained NCC models and apply them to the biological processes mentioned above, see Fig. 6. Our simulation results indicate that the inference approaches explored demonstrate limited success in elucidating the interactions within the examined biological network.

**Figure 6:**
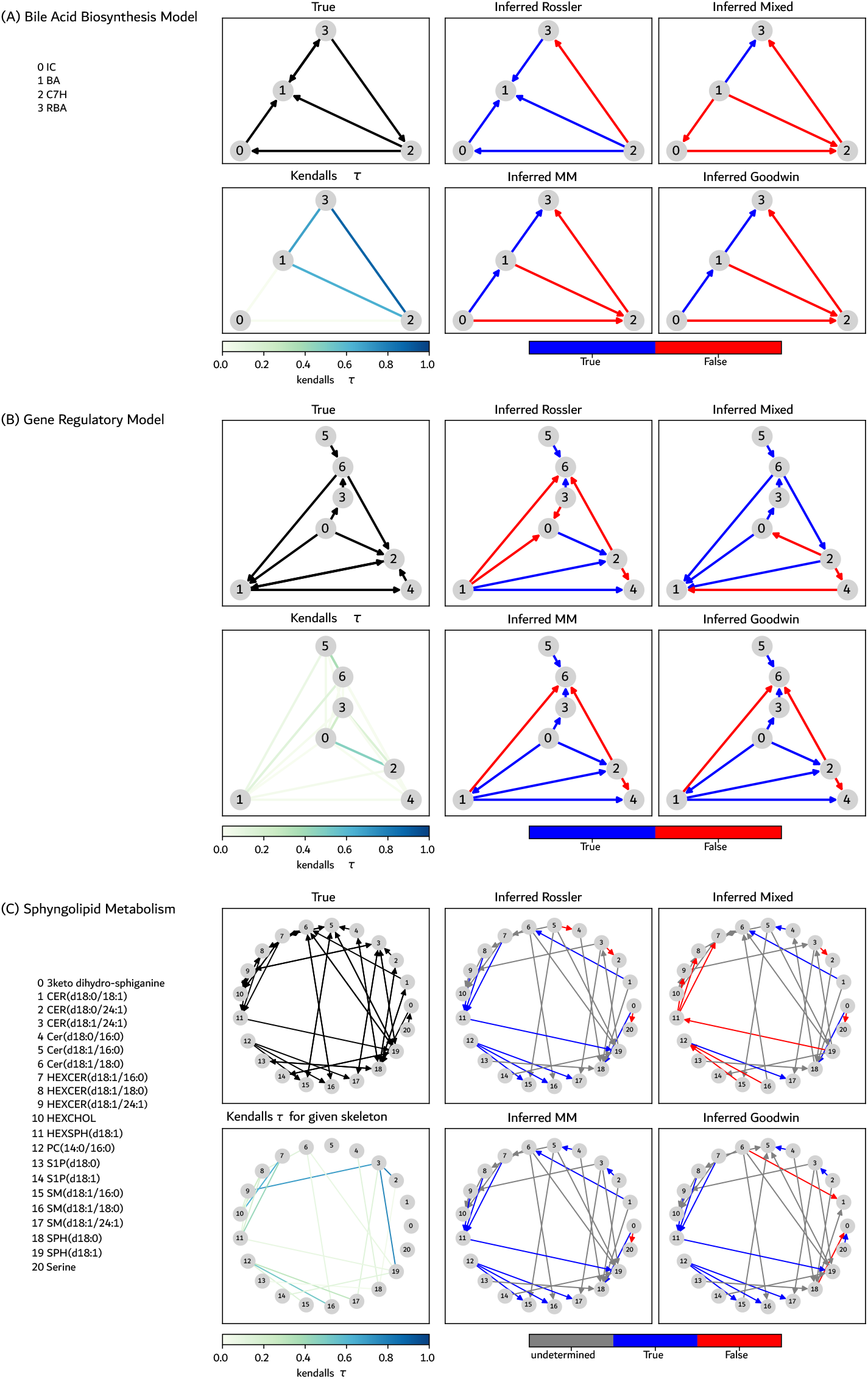
Performance for Realistic Biological Systems. The models used for direction inference are NCC with training data as specified in each panel. Models are detailed in Appendix A

## 3. Discussion and Summary

Inferring the direction of interaction between two components within a complex bio-chemical network from time-independent observations is an important, yet challenging, task. One key significance of inferring directed biochemical networks in human health research lies in the ability to control and recover metabolic performance in faulty or sub-optimally operating cells, such as those with mutations [5]. Additionally, it is essential to understand how signal transduction networks influence cancer cell behavior, including proliferation, survival, invasiveness, and drug resistance[3]. Therefore, the study of biological interaction networks, and in particular the inference of the direction of the biochemical reactions, is essential as it may eventually lead to new therapeutic approaches and personalized treatments.

Here we examined various methods to determine the interaction directionality while considering the characteristics of the biological observations of metabolites; the small data set, many variables that are unknown or unobserved, the multiplicity of confounders, the noise propagation through the network, the unknown dynamics mechanism between the variables, and more. Since many variables are undetermined, we concentrated on pairwise directionality inference.

We compared various approaches and found that they are limited in effectively determining the direction of influence. This limitation may stem from unfulfilled assumptions, such as the presence of directed acyclic graphs or the introduction of additional noise. Furthermore, the complexity can increase due to different interaction mechanisms, multiple undetermined confounders, and various sources of noise. Therefore, we suggest that methods for determining interaction orientation should be specifically tailored to the system being studied whenever possible.

The inference of pairwise directionality from multiple snapshots, especially in the absence of temporal information, should be approached with caution. In particular, one must be wary of interpreting the interaction directionality as indicative of ‘causality.’ This term refers to scenarios where direct changes or interventions in one variable influence the state of another variable. Consequently, ‘causality’ is typically discussed in the context of temporal data, where direct interventions and their subsequent effects can be observed over time. However, while some argue that ‘causality’ should only apply to temporal data, others extend its definition to include an interpretation based on static observations - this dissension remains open for further research.

## Supplementary Materials

## Appendix A. Synthetic Data generation

For numerical demonstrations, we generate a random directed graph *Ĝ* that holds the topology of the network. Mathematically, the graph is described by the adjacency matrix with elements *G*_*ij*_ = 1 where the state of *j* affects the dynamics of node *i*, i.e., *j* → *i*, and *G*_*ij*_ = 0 otherwise. We examined systems with Erdös Rényi random networks similarly to [64].

The synthetic data is generated as follows. First, we generated a long temporal trajectory of (*y*_*i*_, *y*_*j*_) following

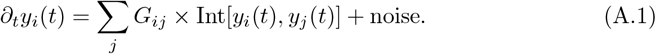

Here *Ĝ* is the adjacency matrix of the Erdös Rényi random interaction network. The function Int[*y*_*i*_, *y*_*j*_] defines both the mechanism of the interaction, the effect of variable *j* ≠ *i* upon the dynamics of variable *i*, and the self-regulation, the effect of the level *y*_*i*_ on oneself dynamics. The noise term can be either additive noise or multiplicative noise.

We note that we do not aim to determine the interaction mechanism Int[*y*_*i*_, *y*_*j*_], and we do not subject ourselves to any specific form of it. Moreover, we note that Int[*y*_*i*_, *y*_*j*_] defines only direct interaction between the variables. However, indirect effects might present as well, especially where the networks we examine might be cyclic graphs due to the nature of the biochemical process.

### Appendix A.1. Models

#### Continuum Michaelis–Menten Regulatory Network

The dynamic in the Michaelis–Menten (MM) regulatory network is given by the

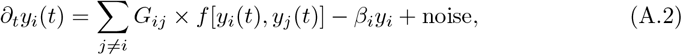

where the interaction function for a given edge is given by

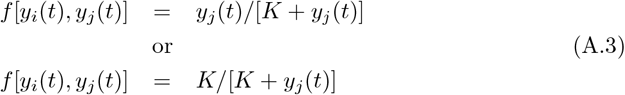

where we choose between the activation (top) and suppression (bottom) randomly, with equal probability to either choice. For variables that are not affected by any other variable, i.e. ∑ _*j*_ *G*_*ij*_ = 0, we use; *f* = 1. The argument of half-maximum rate is determined as *K* = 1. All variables are self-regularized by the term − *β*_*i*_*y*_*i*_, where *β*_*i*_ represents the degradation rate of molecule *i* and is chosen in the simulation to be drawn from Gaussian distribution with mean 1 and noise scale of 0.1.

#### Coupled Goodwin Oscillators

The dynamic for the triplet (*x*_*i*_(*t*), *y*_*i*_(*t*), *z*_*i*_(*t*)) is given by the

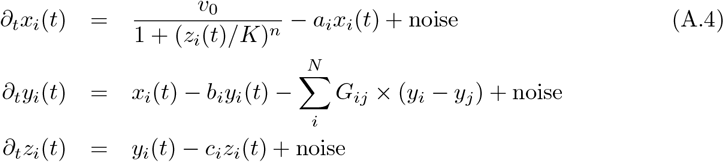

where the numeric simulation took place with the parameters *K* = 1, *a*_*i*_ = *b*_*i*_ = *c*_*i*_ = 0.4, *v*_*o*_ = 1, *n* = 17 as the same as [64].

#### Rössler Oscillators

The dynamic for the triplet (*x*_*i*_(*t*), *y*_*i*_(*t*), *z*_*i*_(*t*)) is given by the

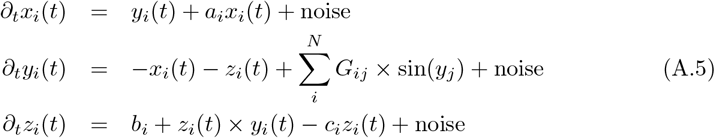

where the numeric simulation took place with the parameters *K* = 1, *a*_*i*_ = 0.1, *b*_*i*_ = 1 and *c*_*i*_ = 18.0 for every *i* as the same as [34].

#### Bile Acid Synthesis Pathway

The Bile Acid Synthesis Pathway (BASP) is modeled by

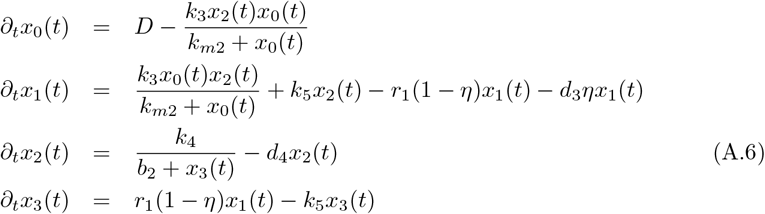

with parameters given in Table below.

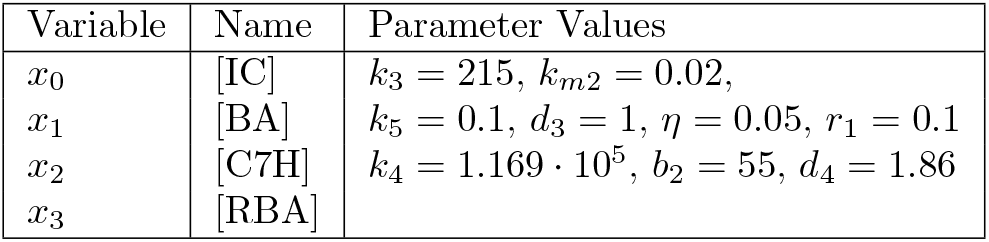

The variables are the concentrations of the intercellular cholesterol [IC], bile acid [BA], the Cholesterol 7*α* Hydroxylase [C7H], and the Returned Bile Acids [RBA]. The ‘external signal’ is modeled by uniform distribution, means *D* ∼ Uniform[0, 1].

#### Gene Regulatory Interactions in E. coli

As mentioned we also examine more biologically realist model which is based on a subset of an *E. coli* gene regulatory network and was provided in the DREAM challenge [57–59]. There, the *in-silico* network inference challenge investigated how well gene networks can be deduced from simulated data. The network is derived as subgraphs from the recognized *E. coli* and *S. cerevisiae* gene regulation networks [72]. Meaning that the results presented are thus biologically realistic, i.e., aiming to capture a reasonable network, but not given from a real observation, and the gene indexes are thus arbitrary. The *E. coli* gene regulatory network is defined and simulated with the following equations:

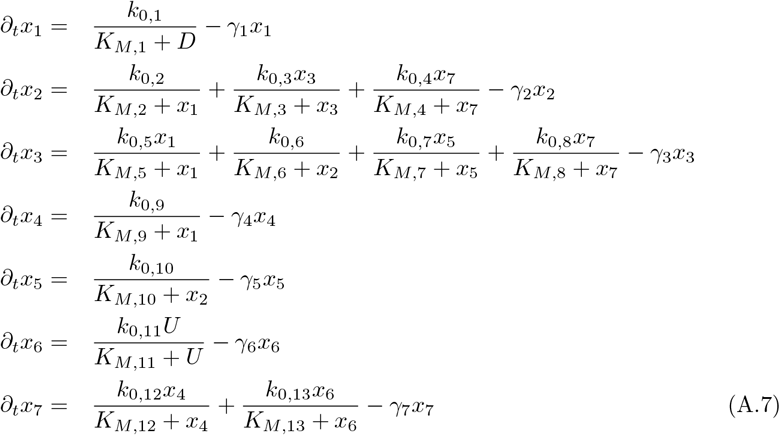

with parameters given in Table below.

| Gene | Parameter Values |
| --- | --- |
| $N_1$ | $k_{0,1} = 0.0362, K_{M,1} = 0.1259, \gamma_1 = 0.4060,$ |
| $N_2$ | $k_{0,2} = 1.0106, K_{M,2} = 1.7937, k_{0,3} = 0.3550,$<br>$K_{M,3} = 1.2069, k_{0,4} = 0.7472, K_{M,4} = 1.2858,$<br>$\gamma_2 = 2.1362$ |
| $N_3$ | $k_{0,5} = 2.4007, K_{M,5} = 0.8218, k_{0,6} = 0.8511,$<br>$K_{M,6} = 1.7099, k_{0,7} = 2.8247, K_{M,7} = 1.6656,$<br>$k_{0,8} = 0.6081, K_{M,8} = 0.0202, \gamma_3 = 3.8740,$ |
| $N_4$ | $k_{0,9} = 0.0903, K_{M,9} = 0.069, \gamma_4 = 0.7256$ |
| $N_5$ | $k_{0,10} = 0.5264, K_{M,10} = 0.9600, \gamma_5 = 0.7466$ |
| $N_6$ | $k_{0,11} = 0.6541, K_{M,11} = 1.0891, \gamma_6 = 0.4525$ |
| $N_7$ | $k_{0,12} = 0.0090, K_{M,12} = 0.5191, k_{0,13} = 1.1236,$<br>$K_{M,13} = 0.4986, \gamma_7 = 0.9473$ |

These parameters are taken from [57–59]. The ‘external disturbance’ is modeled by uniform distribution, means *D, U* ∼ Uniform[0, 1].

#### Sphyngolipid Metabolism

The model includes reactions involved in sphyngomyelin hydrolysis, de novo synthesis of ceramide, and the salvage pathway, where we are specifically following three fatty acid chain molecular species. Specific reactions included in the model, as well as enzymes included in the model, are shown in Figure 6 (C). The abbreviations used in this figure are provided as follows. CER: Ceramide, HEXCER: Hexosylceramide, PC: Phosphatidylcholine, S1P: Sphinganine-1-phosphate, SM: Sphingomyelin, SPH: Sphinganine. The model aims to include the number of enzymes shown to be involved in these reactions (following reaction information obtained from KEGG: map00600, UNIPROT, and Rhea), and does not represent any specific biological situation. The enzyme kinetic is calculated using the method developed by Kroll et al. [73]. The kinetic rate for each enzyme involved in the reaction is calculated separately for specific reactant and product pairs, where the model uses the enzyme sequence and reactant and product SMILES strings to predict the kinetic rate. The mass action model of reaction is calculated using the sum of kinetic parameters for all enzymes involved in the reaction. The model is run until reaching the steady state where the values for serine, 3keto dihydrosphiganine, and PC are kept constant, as necessary metabolites for the system. For other metabolites, we are assuming a closed system. This model was simulated using MATLAB (MathWorks, Inc.). The MATLAB code for this model will be made available upon reasonable request.

## Appendix B. Overview of Pairwise Orientation Methods

Several notable methods have been proposed to address the bivariate causal discovery problem. Early techniques relied on strong assumptions about the dynamic mechanisms, such as additive noise or functional models. However, biochemical variables may not meet these assumptions, particularly when they evolve within complex interaction networks. To determine which methods might be suitable for network orientation analyses, we examine various approaches using synthetic data.

The techniques we examined are based on three features: (1) statistical scores quantified for both directions and presume some statistical dependence (e.g., additional noise model), (2) modeling using neural networks, and (3) an ensemble of models using the concept of meta-learning. Briefly, the statistical scores require us to assume the dynamic mechanism between the two variables but allow inference from a relatively low number of data points. Conversely, methods involving the stacking of statistical features or neural network modeling are heavily data-consuming.

### Appendix B.1. Statistical Scores

Methods for interaction direction inference that involve some statistical scores aim to quantify the asymmetric conditional dependence between the two variables. However, these statistical scores are strongly based on functional causal models (FCM) between the two continuous variables. It means that for two variables *x* and *y*, the edge *x* → *y* means that *y* = *f* (*x, θ, η*) where *θ* is the parameters sets, and *η* is a noise term. The causal structure is identifiable whenever there are no unobserved confounders, it belongs to a restricted functional class, and suitable constraints are imposed on *η*. It has been shown that without any further assumption on the function *f*, the causal direction is not identifiable because for both directions one can find an independent noise term [74, 75] Inspired by [51], we test in the benchmark the following statistical scores.

#### Additive Noise Model (ANM)

The additive noise model (ANM) assumes that *y* = *f* (*x*) + *η* where the noise *η* and variable *x* are independent. The additive noise model is one of the most popular approaches for pairwise causality. It is based on the fitness of the data to the additive noise model in one direction and the rejection of the model in the other direction. The data is assumed to be continuous [61].

#### Conditional Distribution Similarity statistics (CDS)

Assume that the shape of the conditional distribution *p*(*Y* |*X* = *x*) tends to be similar for different values of *x* if the random variable *X* is the cause of *Y* (i.e., if *X* → *Y*). Then, one of the quantities that captures this variability is the standard deviation of the scaled values of y after binning in the x direcLtion. A lower standard deviation indicates *x* → *y*. his measure is defined as: 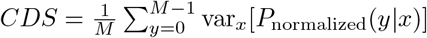.

#### Information Geometric Causal Inference (IGCI)

This approach considers a deterministic process that follows *y* = *f* (*x*), where assumed that the function *f* is invertible, which means that *f* ^−1^ exists. The direction of interaction is then determined by measuring irregularities by the distance to an exponential family using the information statistics metric, i.e., Kullback-Leibler statistics [63]. Furthermore, the authors show the applicability of such statistical measures for processes evolving with small additive noise [63].

#### Regression Error based Causal Inference (RECI)

The approach is based on the assumption that a regression fit in the true causal direction yields smaller errors, on average, than when fitting the model in the opposite direction. It allows non-deterministic nonlinear relations between the cause and the effect. The analysis takes place by fitting a least squares regression in both possible causal directions, and the causal direction is chosen to be the direction with the lower mean-squared errors (MSE) [65]. For the regression implementations, we use polynomial regression of degree 3 [56].

### Appendix B.2. Machine Learning Techniques

Beyond the statistical methods presented above, which, as mentioned, require some strong assumptions or prior knowledge about the causal mechanism, different approaches use machine learning (ML) techniques to detect the direction of the interaction. These methods, however, possess some challenges since they need prior learning of a labeled training set and require large datasets that are not necessarily accessible. Moreover, these methods require additional cost, the need for graphics processing units (GPUs), and computational time due to the complexity of the algorithms.

Generally, ML methods aim to infer the direction of interaction from hidden patterns within the training data, without explicitly coding the features required for inference. It can be done by using neural network (NN) modeling, or by stacking many properties of the system in the so-called meta-learning techniques.

#### Causal Generative Neural Networks (CGNN)

The method aims to learn the multivariate function causal model *f* using a generative NN. The uses of NN to learn *f* allow for not explicitly restricting the class of functions allowed. Particularly, it models *f* for both directions, *y* → *x* and *x* → *y*, with a neural network with one hidden layer. Then it chooses the direction that provides the best fit - using a non-parametric score, the Maximum Mean Discrepancy [67].

#### Randomize Causation Coefficient (RCC)

The method utilizes a NN that is constructed with two parts - kernel embedding layers and a classifier. The former part is based on the projection of the observational distributions into Reproducing Kernel Hilbert Space (RKHS) using random cosine embedding, and the classifier is a random forest [69].

#### Neural Causation Coefficient (NCC)

The neural network (NN) is designed with embedding layers that facilitate the learning of feature maps, which are essential for understanding the data. Following the embedding layers, the architecture includes binary classifier layers, both of which are structured as multilayer perceptrons [68].

#### Stacking with Gradient Boosting Classification - JARFo

This method is an ensemble learning algorithm of many statistical features, including statistical measures, information measures, and measures of the conditional probability variability. The stacking of all these features was done with gradient-boosting classification. In the literature, it is named after its designer - José A. R. Fonollosa (JARFo) - [62].

#### Causal Ensemble Measure Machine (CEMM)

Stacking the statistical scores using the support measure machine (SMM) as follows. First, train SMM, which classifies each score as to whether it successfully detects the direction or not. The stacking involves ‘flipping’ the wrongly assigned direction [51].

